# Intriguing effects of selection intensity on the evolution of prosocial behaviors

**DOI:** 10.1101/2021.03.12.435199

**Authors:** Alex McAvoy, Andrew Rao, Christoph Hauert

## Abstract

In many models of evolving populations, genetic drift has an outsized role relative to natural selection, or vice versa. While there are many scenarios in which one of these two assumptions is reasonable, intermediate balances between these forces are also biologically relevant. In this study, we consider some natural axioms for modeling intermediate selection intensities, and we explore how to quantify the long-term evolutionary dynamics of such a process. To illustrate the sensitivity of evolutionary dynamics to drift and selection, we show that there can be a “sweet spot” for the balance of these two forces, with sufficient noise for rare mutants to become established and sufficient selection to spread. This balance allows prosocial traits to evolve in evolutionary models that were previously thought to be unconducive to the emergence and spread of altruistic behaviors. Furthermore, the effects of selection intensity on long-run evolutionary outcomes in these settings, such as when there is global competition for reproduction, can be highly non-monotonic. Although intermediate selection intensities (neither weak nor strong) are notoriously difficult to study analytically, they are often biologically relevant; and the results we report suggest that they can elicit novel and rich dynamics in the evolution of prosocial behaviors.

**Author summary:** Theoretical models of populations have been useful for assessing when and how traits spread, in large part because they are simple. Rather than being used to reproduce empirical data, these idealized models involve relatively few parameters and are utilized to gain a qualitative understanding of what promotes or suppresses a trait. For prosocial traits, which entail a cost to self to help another, one thing that mathematical models often suggest is that competition to reproduce must be localized, meaning an individual must be fitter than just a small subset of the population in order to produce an offspring. We show here that this finding is not robust. Such traits can indeed proliferate when there is global competition for reproduction, which we demonstrate by increasing the degree to which fitness affects birth rates. Since this kind of “stronger selection” has also been observed empirically, we discuss how it is incorporated into theoretical models more broadly.

## 1 Introduction

The study of prosocial behaviors, which entail personal costs in order to benefit others, has driven the development of many of the models and techniques of modern evolutionary game theory. In its simplest form, evolutionary game theory is a framework for modeling the population dynamics of two or more traits with frequency-dependent reproductive success. The localized interactions between competing types, which influence an individual’s fecundity or survival, are called “games.”

In the donation game, for example, each individual has one of two possible types: *C* (“cooperator/producer”) or *D* (“defector/non-producer”). A producer pays a cost, *c*, to provide the opponent with a benefit, *b*. Non-producers do nothing. Two producers facing one another each receive *b* – *c*, whereas two non-producers each get nothing. But when paired with a producer, an individual is better off not donating, instead taking *b* while the opponent gets – *c*. Although the resulting conflict of interest is quite simple, prisoner’s dilemma interactions like this game have been extremely influential in explaining the evolutionary origins of altruistic behaviors [1].

In order to understand their evolutionary dynamics, these kinds of prosocial traits are studied in conjunction with a rule for updating the population based on reproduction and death. Much of the theoretical work on the evolution of cooperation assumes weak selection, in which payoffs have small effects on reproductive rates. Weak selection is frequently a natural assumption for biological evolution because, in general, an organism with a particular mutation has myriad other traits, and so this one difference between those with wild-type phenotype and the organism with mutated phenotype is not likely to have large effects on overall reproduction rates [2]. Similar remarks can be made about many cultural contexts. In a generic community of people, a single novel behavior might not be significantly noted or adopted, since there are many behaviors exhibited by each person, which may or may not be already shared with others, contributing to the social fabric.

In addition to being a biologically reasonable assumption in many situations, weak selection usually also facilitates the mathematical analysis of evolutionary models. While population structure, which constrains interactions and offspring dispersal, has long been understood (qualitatively) to influence the spread of prosocial behaviors like cooperation [3], more recent studies have sought to uncover quantitative conditions for the spread of cooperation in populations with fine-grained spatial structure. Even in cases for which these conditions ultimately contain terms that can be interpreted as relatedness, the conditions themselves cannot readily be derived just from the intuition that population viscosity increases relatedness between interactants. As such, additional work is required, and weak selection is often a crucial component of such analyses. A well-known study of weak selection is that of death-Birth (*dB*) updating on large regular graphs, which resulted in the widely quoted “*b*/*c* > *k*” rule [4] that says cooperation evolves if the ratio of the benefit provided by a cooperator, *b*, to the cost of cooperation, *c*, exceeds the number of neighbors, *k*. (The capitalization in “death-Birth” refers to the step in which selection acts.) This work has been subsequently generalized in a number of ways, for instance to include distinct interaction and dispersal topologies [5] as well as arbitrary heterogeneous graphs [6].

On the other hand, there are other update rules that seem to never support cooperation, regardless of the population structure. The archetype of such processes is Birth-death (*Bd*) updating, which is inspired by the Moran process [7] and models global competition for reproduction. In well-mixed populations, where the literature historically begins, cooperation in prisoner’s dilemma games is never favored [8]. For *Bd* updating, cooperation is also never favored on regular graphs like the cycle [9], in which every node has the same number of neighbors [4, 10].

The general question of whether prosocial behaviors like cooperation can be favored on any topology, under *Bd* updating and weak selection, is unresolved, but it is generally believed that this is not possible [4, 10, 11]. The intuition behind this is that in *Bd* updating, adjacent individuals, who are playing the prisoner’s dilemma against each other, also compete to reproduce, since *Bd* updating first selects a node to reproduce proportional to fecundity. Then, cooperators are directly helping adjacent and competing defectors to be selected to reproduce. Contrast this to *dB* updating, in which competition to be selected only occurs among the adjacent nodes to the one randomly chosen to die. Two such individuals might themselves not be neighbors, which lessens the chance that an altruist helps a competitor. Therefore, we would expect cooperation to be favored under *dB* updating but not under *Bd* updating [11]. This intuition is captured more broadly and succinctly by Débarre et al. [12], who conclude that “the evolution of social behaviour is determined by the scale at which social interactions affect competition.”

However, weak selection is far from the only relevant paradigm in modeling evolutionary dynamics. There are also compelling examples of strong selection in biology. A classic case study is that of the peppered moth (*Biston betularia*) in England. Prior to the Industrial Revolution, the lighter-colored moths could effectively hide from predators camouflaged amongst lichen and tree bark. Once atmospheric pollution covered the environment in soot, darker peppered moths proliferated (and, in subsequent years, moths began to revert to their original color) [13]. Changing environmental conditions amplified the influence of one particular trait, moth color, above other potential mutations. Rapid temperature changes, e.g. due to climate change, have also been observed to result in increased selection pressures, related to the ability to bear heat, in populations of trees (*Fagus sylvatica*) [14] and wild birds (*Parus major*) [15]. Thus, even for populations in which weak selection might be a viable assumption under normal circumstances, environmental changes can (unsurprisingly) result in non-weak selection for some traits.

Compared with the regime of weak selection, less theoretical work has been done to understand social evolution under intermediate and strong selection, but these areas are not entirely unexplored. “Strong” selection frequently means that the selection intensity parameter is highest, essentially the opposite of weak selection. For prisoner’s dilemma games on cycles, under *Bd* updating and strong selection, it is possible that a community of cooperators can resist invasion and fixation of a mutant defector [16]. More generally, intermediate and strong selection have been studied for multi-player games on the cycle [17], where it is shown that both the multi-player stag hunt game and the snowdrift game can favor cooperation, regardless of the intensity of selection. Theoretical work in this area has also been connected to behavioral experiments, where it is shown that intermediate selection intensities in stochastic models reproduce empirical findings in ultimatum [18] and anticipation [19] games.

In this study, we first discuss several issues that become increasingly important when modeling stronger selection in evolutionary games. These include how to convert payoff to fecundity, how mutants arise within the population, and how to quantify the effects of selection on evolutionary dynamics. We then highlight two important points using examples in simple structured populations. The first demonstrates that models with stochastic payoffs can exhibit completely different dynamics than their deterministic (mean-field) analogues. The second finding is that selection, of the proper intensity, can favor the evolution of prosocial behaviors under *Bd* updating, which challenges the consensus that *Bd* updating is necessarily a hostile setting for altruism. In particular, prosocial behaviors can evolve even when there is global competition for reproduction.

## 2 Results

### 2.1 Evolutionary games and the intensity of selection

When selection is neither weak nor strong, three concerns immediately arise regarding how to model and evaluate evolutionary dynamics. The first issue relates to how payoffs accrued in games are converted to fecundity. A number of functional forms have been proposed in the literature, but all of them are effectively equivalent under weak selection, so there is no need to distinguish between different payoff-to-fecundity maps in that case. When selection is not weak, we show that there is a unique payoff-to-fecundity map, provided some natural qualitative assumptions hold.

The second concern is with how mutants arise in the population. Here, we focus on processes in which there is either no mutation or for which mutation is sufficiently rare. As a result, once a mutant does appear, the evolutionary process will drive the mutant type either to extinction or fixation prior to the arrival of another mutant. This leads to a useful metric for characterizing the evolutionary success of the mutant type: its fixation probability, which is the probability that the mutant type eventually takes over the population prior to the occurrence of another mutation. While different kinds of mutant appearance distributions are often considered in the literature on weak selection, here the issue becomes a bit more nuanced.

Finally, for processes with sufficiently rare mutation, we arrive at the question of how to quantify the evolutionary success of a trait. Under weak selection, this question is typically approached in one of two ways, either absolutely (“does selection increase the fixation probability of cooperators?”) or relatively (“is the fixation probability of cooperators greater than that of defectors?”). Versions of these two conditions are still valid and reasonable for non-weak selection, but they are subtler. Under weak selection, the reference point is neutral drift (by definition), whereas for stronger selection intensity, represented by some parameter *δ*, one could choose either neutral drift or *δ* as a reference point. The absolute condition bifurcates into two conditions: “is the fixation probability at *δ* greater than that of neutral drift?” and “does increasing selection slightly beyond *δ* increase the fixation probability, compared to its value at *δ*?” When *δ* is sufficiently close to neutral drift, these two conditions collapse into one. There is a similar bifurcation for the relative condition, which gives a total of four ways in which we quantify the evolutionary success of a trait when selection is not necessarily weak: two absolute conditions and two relative conditions.

#### 2.1.1 Payoff accrual

Here, we consider payoffs arising from simple prosocial behaviors. In particular, matrix games among two players do not specify how individuals accrue payoffs through interactions in a population. Naturally, this represents a prerequisite for understanding and analyzing the resulting evolutionary dynamics.

In fact, there is no canonical way to define such a quantity, particularly when the population structure is heterogeneous. Two of the most popular ways of defining an individual’s overall payoff are through accumulation and averaging. In the former, an individual simply adds together the payoffs from all pairwise encounters, while in the latter these payoffs are averaged. Both methods involve deterministic payoffs, in which each individual receives an amount that is uniquely specified by the population’s structure and configuration of producers and non-producers. However, both kinds of payoff aggregation in a two-player game are just the beginning of the story of prosocial traits in structured populations. More specifically, the donation game is additive, which means that an individual’s payoff in an interaction can be decomposed into the payoff from his or her action plus the payoff from the action of the opponent. Therefore, rather than thinking of payoffs as the result of pairwise *interactions*, in the donation game it is also natural to think of payoffs arising from a sequence of individual *actions*, one for each individual in the population. Incidentally, this action-based interpretation is why we use the terms “producer” and “non-producer” in place of the more traditional labels “cooperator” and “defector.”

This interpretation naturally gives rise to other kinds of payoff accounting, such as concentrated or diffuse benefits [20]. When benefits are concentrated, every producer chooses a random neighbor and pays a cost, *c*, to confer a benefit, *b*, on this neighbor only (“*cf*-goods,” where “*c*” indicates that the benefit is concentrated and “*f*” indicates that the cost is fixed). No action is taken to benefit other neighbors. When benefits are diffuse, each producer pays the same cost, *c*, but divides the benefit equally among all neighbors so that each neighbor gets at least a small benefit (“*ff*-goods” since both the total benefit and the total cost are fixed, irrespective of the number of neighbors a producer has). The latter may be viewed as a mean-field version of the former since, under concentrated benefits, the expected payoff to any given neighbor is exactly what this neighbor would receive under diffuse benefits. The classical setting is that of “*pp*-goods,” wherein the total benefit and the total cost are proportional to the number of neighbors a producer has. Depictions of these traits, together with the classical payoff scheme, are shown in **Fig. 1**. The structures considered here, the cycle (**Fig. 1**(*a*)) and the star (**Fig. 1**(*b*)), represent two extremes with regards to payoff accrual. In the former, payoffs are distributed in the same way throughout the population; in the latter, there is a single individual whose connections vastly outnumber those of others.

**Figure 1:**
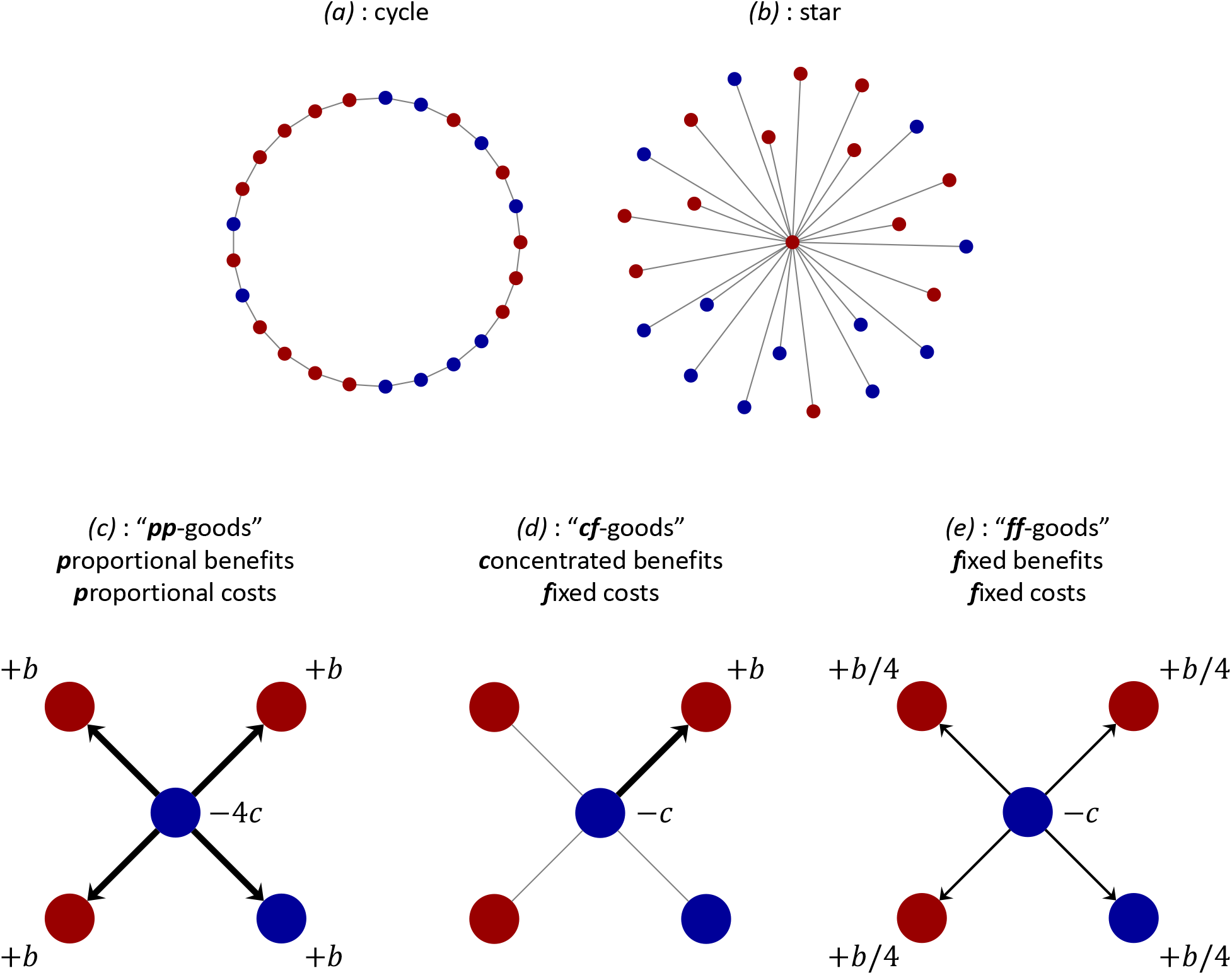
(*a*,*b*) Two examples of spatial structure in a population of size *N* = 25. Each node indicates an individual, who is either a producer (blue) or non-producer (red). Links between nodes represent possible interaction partners and dispersal patterns. The graph depicted in (*a*) is a cycle, which is a periodic structure in which every individual has exactly two neighbors. The graph in (*b*) is a star, a heterogeneous structure consisting of one central “hub” and N – 1 “leaf” nodes at the periphery. (*c*,*d*,*e*) Prosocial behaviors derived from the production of social goods. (*c*) represents the typical model, wherein each producer donates b to each neighbor, at a per-neighbor cost of *c*. In (*d*), every producer confers the entire benefit produced, *b*, on a single, randomly-chosen neighbor. In (*e*), this benefit is divided up evenly among all *k* neighbors. In both (*d*) and (*e*), every producer in the population generates a total benefit of *b* and pays a total cost of *c*; the difference between the two is the way in which this benefit is distributed. Payoffs are a stochastic function of state in (*d*) and a deterministic function of state in (*e*). Whereas (*c*) is *interaction-based*, the models of concentrated and diffuse benefits in (*d*) and (*e*) are more naturally seen as action-based payoff specifications.

#### 2.1.2 Converting payoff to fecundity

In evolutionary games, the payoff, *u*, to an individual is converted to relative fecundity through a payoff-to-fecundity map, *F*(*u*). We impose the following natural and intuitive restrictions on this map. (*i*) In a mechanistic approach to evolutionary dynamics, birth and death should be treated separately rather than amalgamating them into one quantity known as “fitness” [21]. Each of these separate components make sense only when non-negative; and as such, the propensity to reproduce, i.e. fecundity, should also be non-negative. (*ii*) Increasing payoffs are associated to greater abilities in competition, and hence *F* should be non-decreasing. (*iii*) Continuity of *F* is natural in the sense that it implies sufficiently small changes in payoffs do not result in large jumps in fecundity. (*iv*) Selection intensity is interpreted as an amplification of the payoffs prior to the conversion to fecundity. (*v*) Finally, competition should be independent of any baseline payoff or wealth shared by everyone. In particular, this last assumption allows one to focus on the effects of a particular game on competition to reproduce. Thus, in slightly more detail, we have the following assumptions:

i. *F* (*u*) ⩾ 0 for every 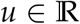;
ii. *F* is non-decreasing;
iii. *F* is continuous;
iv. selection intensity scales payoffs, i.e. the fecundity of payoff *u* at intensity *δ* ⩾ 0 is *F* (*δu*);
v. for every competition within a group *I* ⊆ {1,…, *N*}, the probability that *i* ∈ *I* is chosen for reproduction is invariant under adding a constant, *K*, to the payoffs of all competing individuals in *I*. That is, if *u_i_* and *F_i_* are the payoff and fecundity of individual *i*, then

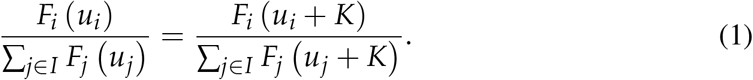

In Methods, we show that, up to a rescaling of the selection intensity, these assumptions lead to a unique payoff-to-fecundity map, *F* (*u*) = *e^δu^* for selection intensity *δ*. Exponential payoff-to-fecundity maps like this one are already commonly used in the literature [17, 22–24], although often with little discussion for the choice of this particular functional form. One argument is that it satisfies assumption (*i*) for all possible payoffs, *u*, precluding complications arising from functional forms like *F* (*u*) = 1 + *δu* [10, 25] or *F* (*u*) = 1 – *δ* + *δu* [8], which can both be negative when *u* is sufficiently small. Under weak selection (*δ* ≪ 1), these functional forms are all equivalent, so one can simply make an arbitrary choice. Another argument is that this kind of exponential mapping matches imitation processes inspired by statistical mechanics, such as a standard model of pairwise comparison dynamics [26, 27]. Our contribution is to realize this common mapping as the consequence of several natural assumptions on the effects of payoffs on competition.

If selection acts on survival rather than on reproduction, we obtain a similar result. However, assumptions (*ii*) and (*v*) need to be modified accordingly. We discuss selection acting on survival further in Methods.

#### 2.1.3 Fixation probabilities and mutant initialization

A standard method for determining the effects of selection on a trait is through fixation probability [8, 28]. Let *ρC.i* (resp. *ρ_D,i_*) denote the probability that a single producer (resp. non-producer) at location *i* generates a lineage that takes over the population. Suppose that *μ_C,i_* (resp. *μ_D,i_*) is the probability that a mutant *C* (resp. *D*) appears at location *i* in the monomorphic all-nonproducer (resp. all-producer) state. The mean fixation probabilities of the two types are then, by definition, 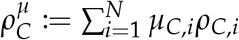 and 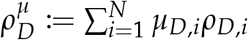.

Under uniform initialization, the probability that an invading mutant (of either type) appears at location *i* is 1/ *N*. Uniform mutant appearance is relevant, for example, when mutations are not tied to reproduction (e.g. external mutagens such as cosmic rays, or the exploration of new behaviors or ideas), or when death rates are roughly the same throughout the population. We denote the mutant-appearance distribution, *μ*, under uniform initialization by *μ*^unif^, which satisfies 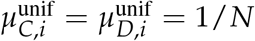. In this case, *μ*^unif^ is independent of the selection intensity, *δ*. We denote by 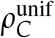 and 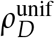 the corresponding mean fixation probabilities.

Another natural distribution is based on mutations that arise during reproduction, and it depends on selection intensity since these mutations are influenced by the rates at which nodes are replaced. In the monomorphic all-producer (resp. all-non-producer) state, let *d_C,i_* (resp. *d_D,i_*) be the probability that the individual at location *i* is replaced by the offspring of another individual, in one step of the process. Since mutations arise during reproduction, higher replacement rates naturally correspond to greater probabilities of mutant appearance. In the limit of rare mutation, we obtain “temperature” initialization, in which the probability that a mutant arises at location *i* is 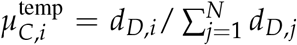 (resp. 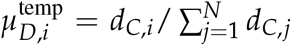) following the all-non-producer (resp. all-producer) state. We denote by 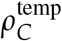 and 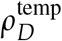 the mean fixation probabilities under temperature initialization.

Notably, uniform and temperature initialization need not coincide. As an example, we consider Birth-death updating, in which an individual is first selected to reproduce with probability proportional to fecundity, and then the offspring replaces a random neighbor. Let 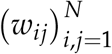 be the adjacency matrix for the population structure, and let 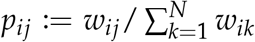 be the probability of moving from *i* to *j* in one step of a random walk on the graph. If *F_i_* denotes the fecundity of individual *i* at a given point in time, then the probability that node *i* is replaced is

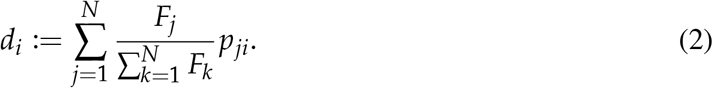

Since 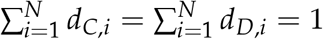, we have 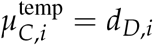 and 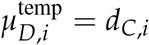.

Consider the star graph (**Fig. 1**(*b*)). In the monomorphic non-producer state, everyone has the same payoff, regardless of selection intensity. As a result, the probability that the hub reproduces is 1/*N*, and the probability that a leaf node reproduces is 1 – 1/*N*. This means that a mutant arises at a leaf with probability 1/*N* and at the hub with probability 1 – 1/*N*. In the monomorphic producer state, reproduction rates depend on the selection intensity. When *δ* ≪ 1, a mutant arises at a leaf with probability approximately 1/*N* and at the hub with probability approximately 1 – 1/*N*. But as *δ* grows, the former probability approaches 1, while the latter approaches 0. **Fig. 2** illustrates temperature initialization on the star in these two states for producers of *ff*-goods (**Fig. 1**(*e*)) under Birth-death updating.

**Figure 2:**
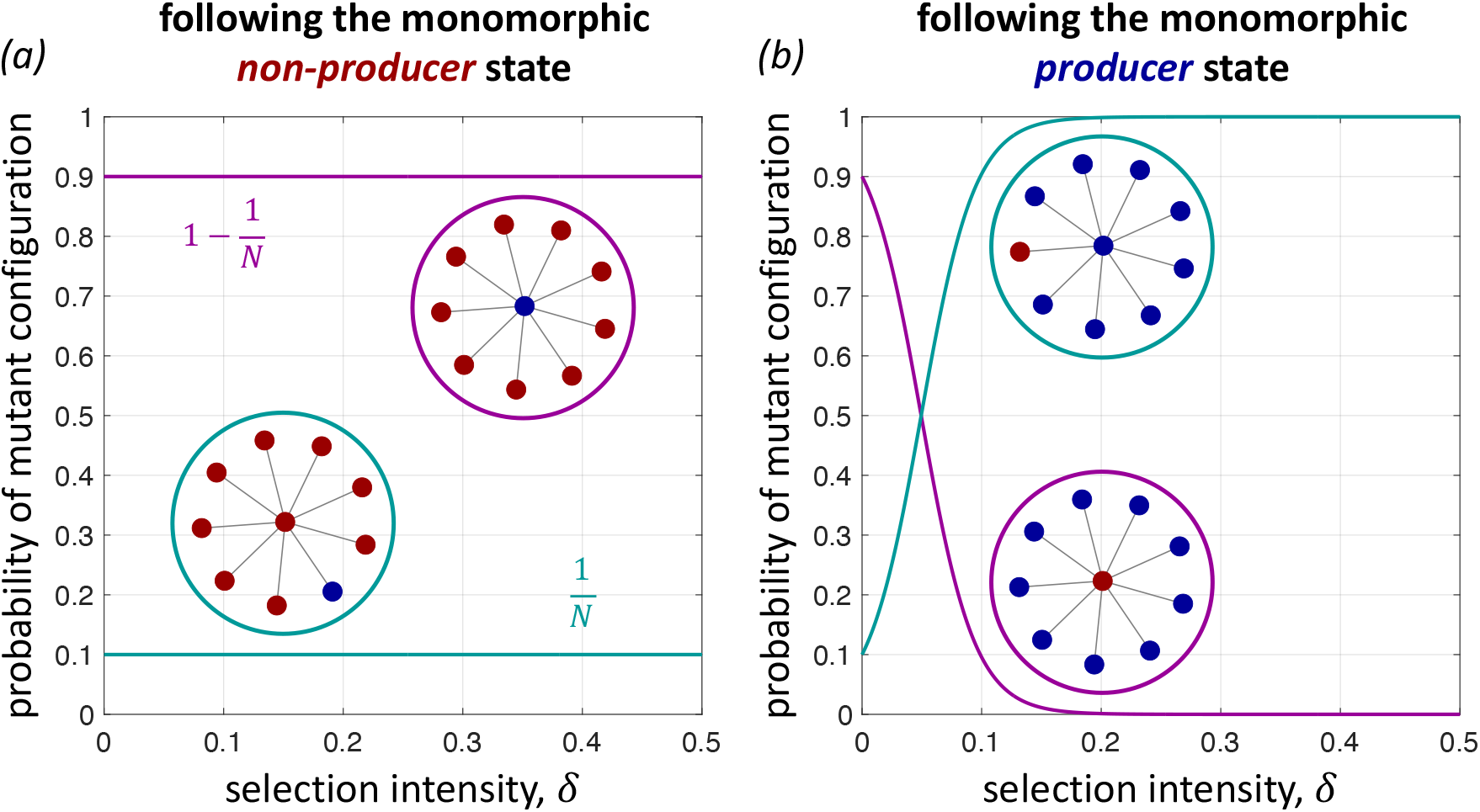
Temperature-based mutant initialization for *ff*-goods under Birth-death updating. (*a*) Following the monomorphic non-producer state, a mutant producer appears either in the hub (purple) or a leaf (turquoise). Since everyone has the same payoff (zero) in the all-non-producer state, the death rates are the same as those of neutral drift, for all *δ* ⩾ 0. With probability 1/*N*, the hub gives birth and the mutant offspring is propagated to a leaf node. Otherwise, with probability 1 – 1/*N*, a leaf node gives birth and the mutant arises in the hub. (*b*) In contrast, death rates in the monomorphic producer state depend on selection intensity. Outside of weak selection, the hub is overwhelmingly more likely to reproduce than a leaf node, meaning a mutant appears in a leaf (turquoise). Although a non-producer at the hub, surrounded by producers, would have a high payoff and thus a high probability of spreading its behavior throughout the population, this mutant configuration appears with probability rapidly approaching zero as *δ* grows (purple). Parameters: *N* = 10, *b* = 5, and *c* = 1.

#### 2.1.4 Quantifying the effects of selection

We now propose four natural ways to quantify selection favoring *C*:

(*A*^0^) *ρ_C_* (*δ*) > *ρ_C_* (0) (absolute; in comparison to neutral drift);
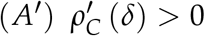 (absolute; in comparison to selection intensity *δ*);
(*R*^0^) *ρ_C_* (*δ*) > *ρ_D_* (*δ*) (relative to *D*; in comparison to neutral drift);
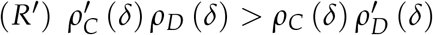 (relative to *D*; in comparison to selection intensity *δ*).

Condition (*A*^0^) disregards the fixation probability of non-producers and requires selection to increase the fixation probability of producers over its value at neutral drift. The metric is “absolute” in the sense that it does not take into account the fixation probability of *D*. Outside of weak selection, this condition involves the comparison of two processes that are quite different: one in which there are no reproductive differences between traits and one in which these differences are substantial.

Condition (*A*′) states that, at a selection intensity of *δ*, increasing the selection intensity beyond *δ* tends to improve the chances that a rare producer can invade and take over a population of nonproducers. This condition is “absolute” for the same reason (*A*^0^) is, but the point of comparison is the fixation probability of *C* at *δ*, and *δ* is not necessarily small.

Condition (*R*^0^) says that *C* is favored relative to *D* if the probability that a rare producer takes over a population of non-producers exceeds the probability that a rare non-producer takes over a population of producers. Equivalently, this condition says that

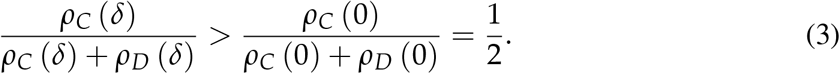

In a process with sufficiently rare mutation, the quotient *ρ_C_*/ (*ρ_C_* + *ρ_D_*) represents the amount of time spent in the all-producer state; the remaining time is spent in the all-non-producer state [29]. Thus, condition (*R*^0^) is a measure of *C* relative to *D*, and its point of comparison is neutral drift. If this condition holds, then at a selection intensity of *δ*, the population spends a larger fraction of its time in the all-producer state than it does under neutral drift. This kind of measure, based on invasion and counter-invasion, is a natural way to quantify evolutionary stability in finite populations [30, 31].

Finally, condition (*R*′) is equivalent to (*ρ_C_* (*δ*) / (*ρ_C_* (*δ*) + *ρ_D_* (*δ*)))′ > 0. It says that, at a selection intensity of *δ*, increasing the selection intensity tends to increase the fraction of time the population spends in the all-producer state. Thus, it provides the same kind of relative measure as (*R*^0^), but the point of comparison is *δ*.

In the limit of weak selection, *δ* ≪ 1, conditions (*A*^0^) and (*A*′) are equivalent, as are conditions (*R*^0^) and (*R*′), and both of these sets of equivalent conditions have been studied. Conditions (*A*^0^) and (*A*′) can be equivalent to (*R*^0^) and (*R*′) under weak selection, for example in games on graphs with the “equal gains from switching” property such as the donation game [4, 8], but they need not always be [see 32]. For stronger selection, all four conditions are generally distinct, and (to our knowledge) they have not been considered individually. In the subsequent examples, we consider these conditions as a way to classify and understand the effects of selection on the evolution of producers of social goods.

### 2.2 Moderate selection intensities in heterogeneous structures

We begin with an example of death-Birth (*dB*) updating, an evolutionary update rule known to support the evolution of prosocial behaviors in structured populations [4, 33]. Under *dB* updating, an individual is first selected for death uniformly-at-random from the population. The neighbors of this individual then compete to fill the vacancy with probability proportional to relative fecundity. A neighbor’s offspring inherits the parental type (either *C* or *D*), replaces the deceased individual, and the process repeats. *dB* updating is homogeneous in the sense that the individual chosen for replacement is always chosen uniformly, regardless of the population’s structure or selection intensity. In particular, uniform and temperature initialization coincide, so there is no need to consider the two mutant-appearance distributions separately.

**Fig. 3** depicts evolutionary dynamics of three kinds of producers of social goods on the cycle (**Fig. 1**(*a*)). Due to the homogeneity of the cycle, all initially-rare mutant configurations are equivalent. Since every individual on the cycle has exactly two neighbors, the difference between *pp*-goods (**Fig. 1**(*c*)), those of benefit and cost proportional to the number of neighbors, and *ff*-goods (**Fig. 1**(*e*)), those of fixed benefit and cost, amounts to a rescaling of the selection intensity. Hence, **Fig. 3**(*c*) appears as simply a horizontally-stretched version of **Fig. 3**(*a*).

**Figure 3:**
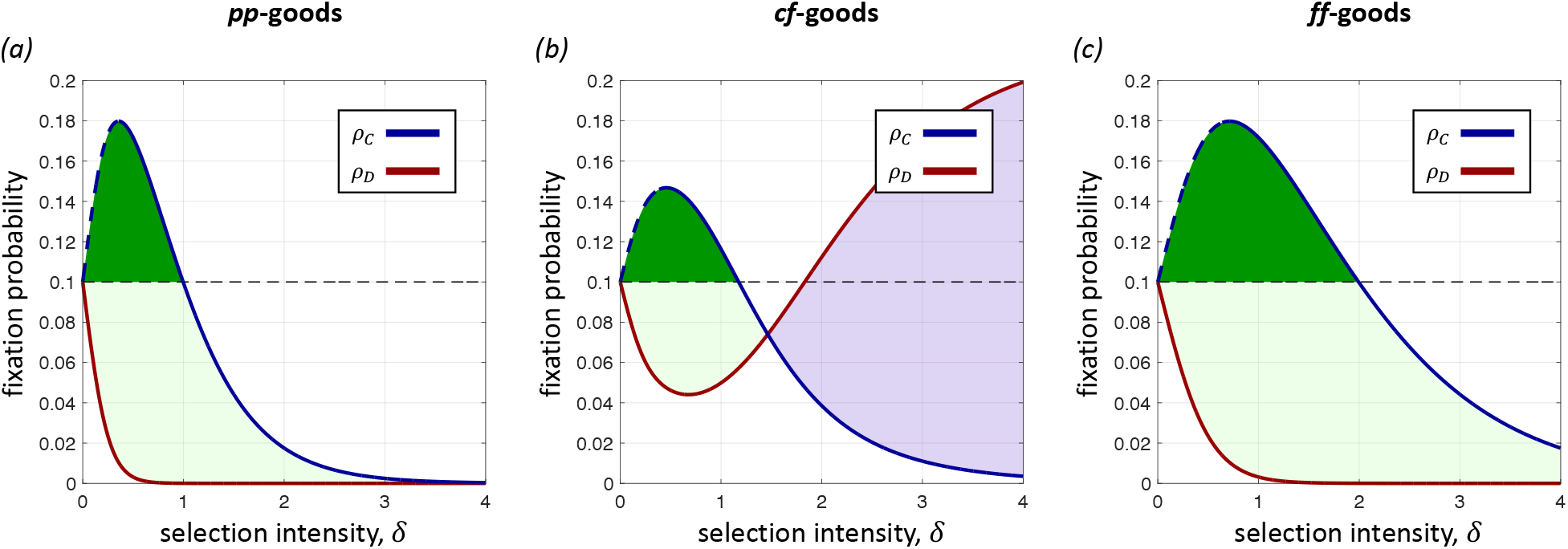
Fixation probabilities with uniform initialization (= temperature initialization) on the cycle of size *N* = 10 under *dB* updating (exact calculations). Since the cycle is a regular graph, the difference between *pp*-goods and *ff*-fgoods amounts to a scaling of the selection intensity. In both cases, (*a*) and (*c*), *C* is always favored by selection relative to *D* (dark+light green), i.e. condition (*R*^0^). But in (*b*), when benefits are concentrated, *ρ_D_* eventually exceeds *ρ_C_*; beyond this intensity of selection, *C* is disfavored relative to *D* (purple, condition (*R*^0^)). Notably, non-weak selection highlights significant qualitative differences between the stochastic payoff scheme of (*b*) and its mean-field analogue, (*c*). In all three cases, *C* is favored absolutely for small *δ*, both with respect to neutral drift (dark green, condition (*A*^0^)) and with respect to *δ* (blue dashed line, condition (*A*′)). The fixation probability under neutral drift is indicated by a black dashed line. For simplicity, we do not illustrate condition (*R*′). Parameters: *b* = 4 and *c* = 1.

What is more notable is the fact that *cf*-goods (**Fig. 3**(*b*)), those of concentrated benefit and fixed cost, and its mean-field analogue, *ff*-goods (**Fig. 3**(*c*)), have completely different behavior apart from when the selection parameter is small. Suppose that a non-producer arises in a population of producers of *ff*-goods. In order for non-producers to fix, the population must transition through a state containing a cluster of exactly two non-producers. In order to acquire additional non-producers from this state, a producer on the boundary must be selected for death. The nonproducer competing to fill the vacancy has a payoff of *b*/2, while the producer on other side of the deceased has a payoff of *b* – *c*. As a result, the probability of acquiring an additional non-producer in this state is (2/*N*) (*e*^*δb*/2^/ (*e*^*δb*/2^ + *e*^*δ*(*b*–*c*)^)), where the factor of 2 is due to there being two producer/non-producer boundaries. Provided *b* > 2*c*, this probability approaches zero as *δ* grows, which effectively precludes non-producers from replacing producers. But consider what happens in this state for producers of cf-goods. If the competing non-producer gets *b*, then the competing producer necessarily has at most *b* – *c* because the shared producer, who died, can donate to exactly one of these two individuals. Thus, the probability of acquiring an additional non-producer does not vanish in this state as *δ* grows. Similar reasoning in the other transient states explains the dynamics of non-producers depicted in **Fig. 3**(*b*).

The similarities between all three panels in **Fig. 3** when *δ* is sufficiently small is no coincidence. In the limit of weak selection, it is known that one can replace stochastic payoffs with deterministic payoffs (i.e. expected payoffs) without a loss of generality [20]. In particular, *cf*- and *ff*-goods exhibit identical dynamics when selection is sufficiently weak, on any population structure. **Fig. 3** demonstrates that this finding does not extend to stronger selection intensities. If one were to model stochastic payoffs using an expected-payoff approach, the model would show that stronger selection favors producers relative to non-producers when, in actuality, that is not the case (**Fig. 3**(*b*)). A description of how the exact numerical calculations in **Fig. 3** were carried out may be found in Methods.

While *dB* updating highlights important differences between the dynamics stochastic and deterministic payoffs, much more curious phenomena arise under Birth-death (*Bd*) updating. Under *Bd* updating, in each time step an individual is chosen for reproduction with probability proportional to relative fecundity. The clonal offspring then replaces a random neighbor. One major difference between *dB* and *Bd* updating is that fecundity acts locally in the former and globally in the latter. As a result, *Bd* updating is not homogeneous when selection acts on the traits; differential birth rates result in neighbors of highly successful individuals being replaced more than those with lower fecundity.

For both kinds of mutant initialization, uniform and temperature, **Fig. 4** depicts evolutionary dynamics of three kinds of social goods on the star (**Fig. 1**(*b*)). For *pp*-goods, producers are disfavored, both absolutely and relative to non-producers, for all *δ* when mutant initialization is uniform. When mutant initialization is temperature-based, producers are still disfavored absolutely, but they are favored relative to non-producers provided *δ* is sufficiently large. The reason for this difference is precisely the behavior described in **Fig. 2**(*b*), wherein the only state with a rare nonproducer becomes increasingly unlikely to appear as an initial condition. We note that under weak selection, the behavior depicted in all panels of **Fig. 4** is essentially the same, with producers disfavored both absolutely and relative to *D*.

**Figure 4:**
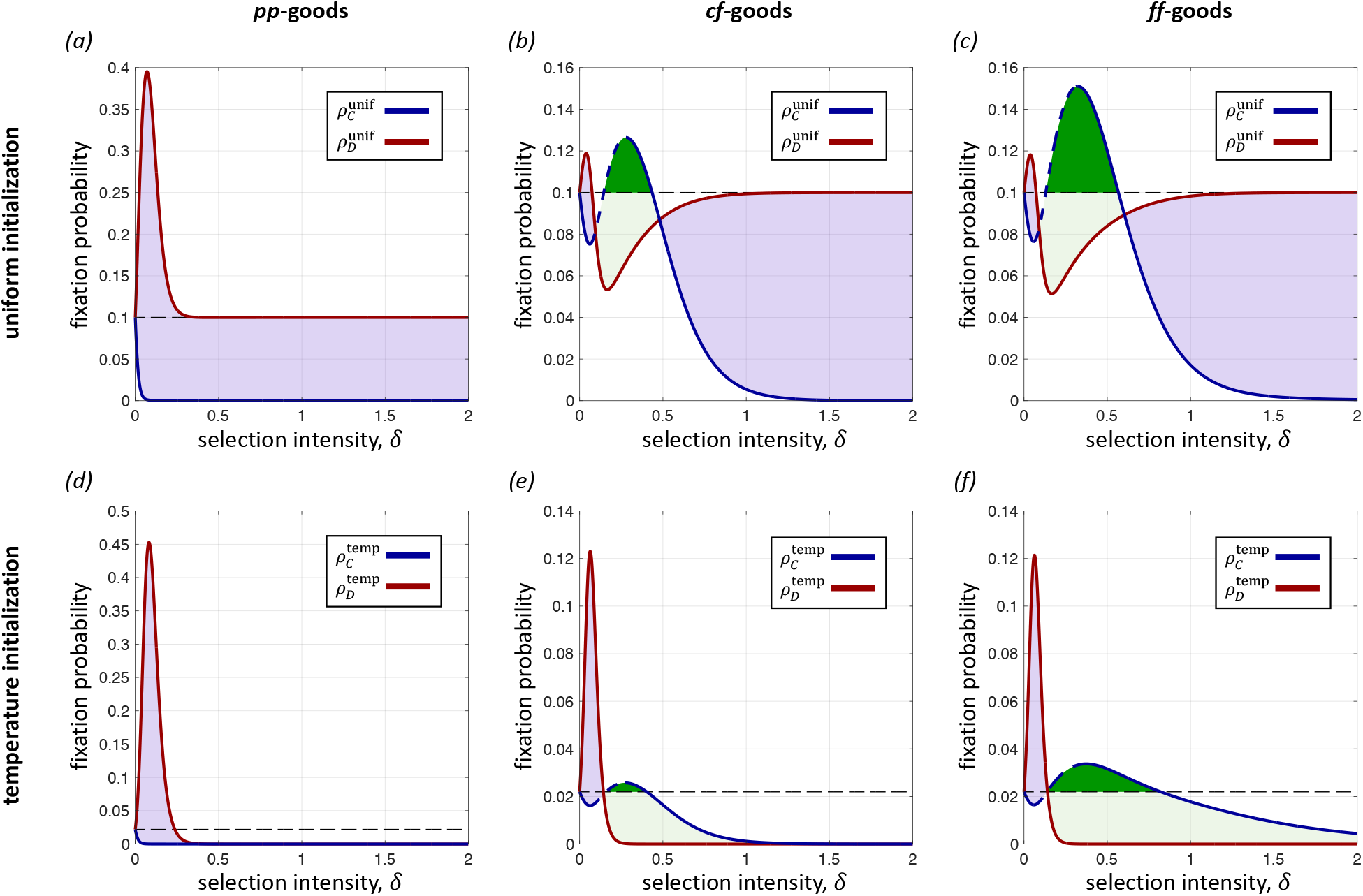
Fixation probabilities on the star of size *N* = 10 under *Bd* updating (exact calculations). (*a*,*b*,*c*) depict the dynamics of social goods under uniform initialization, whereas (*d*,*e*,*f*) show those of death-rate-based temperature initialization. In each panel, we see that weak selection disfavors producers, consistent with earlier studies. However, for both kinds of mutant initialization, we observe the remarkable fact that producers of *cf*- and *ff*-goods can be favored, both absolutely (dark: green, condition (*A*^0^); blue dashed line, condition (*A*′)) and relative to non-producers (dark+light green, condition (*R*^0^)), for a range of intermediate selection intensities. This range represents a “sweet spot” for the population, wherein drift is strong enough to allow for configurations favorable to producers to get established, and selection is strong enough to push those favored configurations toward fixation. The purple regions represent selection intensities for which *C* is disfavored relative to *D* (condition (*R*^0^)). Black dashed lines indicate fixation probabilities under neutral drift. For simplicity, we do not show condition (*R*′). Parameters: *b* = 5 and *c* = 1.

More fascinating is the fact that under both *cf*- and *ff*-goods, for both kinds of mutant initialization, producers can be favored both absolutely and relative to non-producers, provided selection is neither too weak nor too strong (**Fig. 4**(*b*,*c*,*e*,*f*)). Weak selection initially favors non-producers and disfavors producers, consistent with traditional observations about *Bd* updating. However, past a certain selection intensity, fixation probabilities of producers start to increase, even going significantly above their respective values at neutrality. Those of non-producers, in contrast, decline and then dip below their respective values at neutrality. Once *δ* becomes sufficiently large, the fixation probabilities of producers then fall below their neutral values once again. In the case of uniform initialization, the fixation probability of non-producers eventually rises above that of producers (**Fig. 4**(*b*,*c*)).

What is happening at these intermediate selection intensities? For one, drift is still strong enough that the population can transition into a favorable state for producers, such as the one with a producer at the hub and a small number of producers at the periphery. At this point, using *ff*-goods as an example, the producer at the hub has a greater payoff than all other individuals in the population because they pay a single cost, *c*, but receive the whole benefit, *b*, from each of several neighbors. While drift is strong enough to visit such a state, so too is selection strong enough to drive the population to an all-producer state. The individual at the hub is chosen for reproduction disproportionately often, and the offspring must replace individuals at the periphery, eventually driving non-producers to extinction. When selection is too weak, these favorable states are not favored enough to overcome the effects of drift. When selection is too strong, these favorable states are unlikely to be visited in the first place.

For *pp*-goods, we do not observe the same dynamics at intermediate selection intensities. Once a mutant producer occupies the hub, it is comparatively more difficult for producers to replace nonproducers. In contrast to *ff*-goods, there are additional costs to a producer of *pp*-goods at the hub since it must pay a separate cost for each neighbor. Moreover, *pp*-goods offer additional benefits for those at the leaves since the central producer provides each neighbor with a full (rather than fractional) benefit. We note that, due to the rapid decay of mutant appearance at the hub in the all-producer state (**Fig. 2**(*b*)) under temperature initialization, producers are eventually favored relative to non-producers, even though neither one is favored absolutely at strong selection intensities (**Fig. 4**(*d*)).

Continuing with *Bd* updating on the star graph, we also consider significantly larger population sizes. We focus on producers of *ff*-goods because (*i*) at intermediate selection intensities, their dynamics are significantly more interesting than those of producers of *pp*-goods and (*ii*) on the star, there is little qualitative distinction between the dynamics of producers of *ff*- and *cf*-goods. **Fig. 5** depicts the fixation probabilities of producers of *ff*-goods under temperature initialization for population sizes ranging from *N* = 10 to *N* = 2000, extending the example of **Fig. 4**(*f*). At its peak, strikingly, the fixation probability of a single producer is nearly 140-fold its neutral value when *N* = 2000 (**Fig. 5**(*a*)). Further nonlinearities also arise as the population size increases. For sufficiently large *N*, 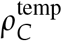 has two distinct local maxima as a function of *δ*, the first of which being attributed to the leaf nodes and the second being attributed to the hub (**Fig. 5**(*b*)).

**Figure 5:**
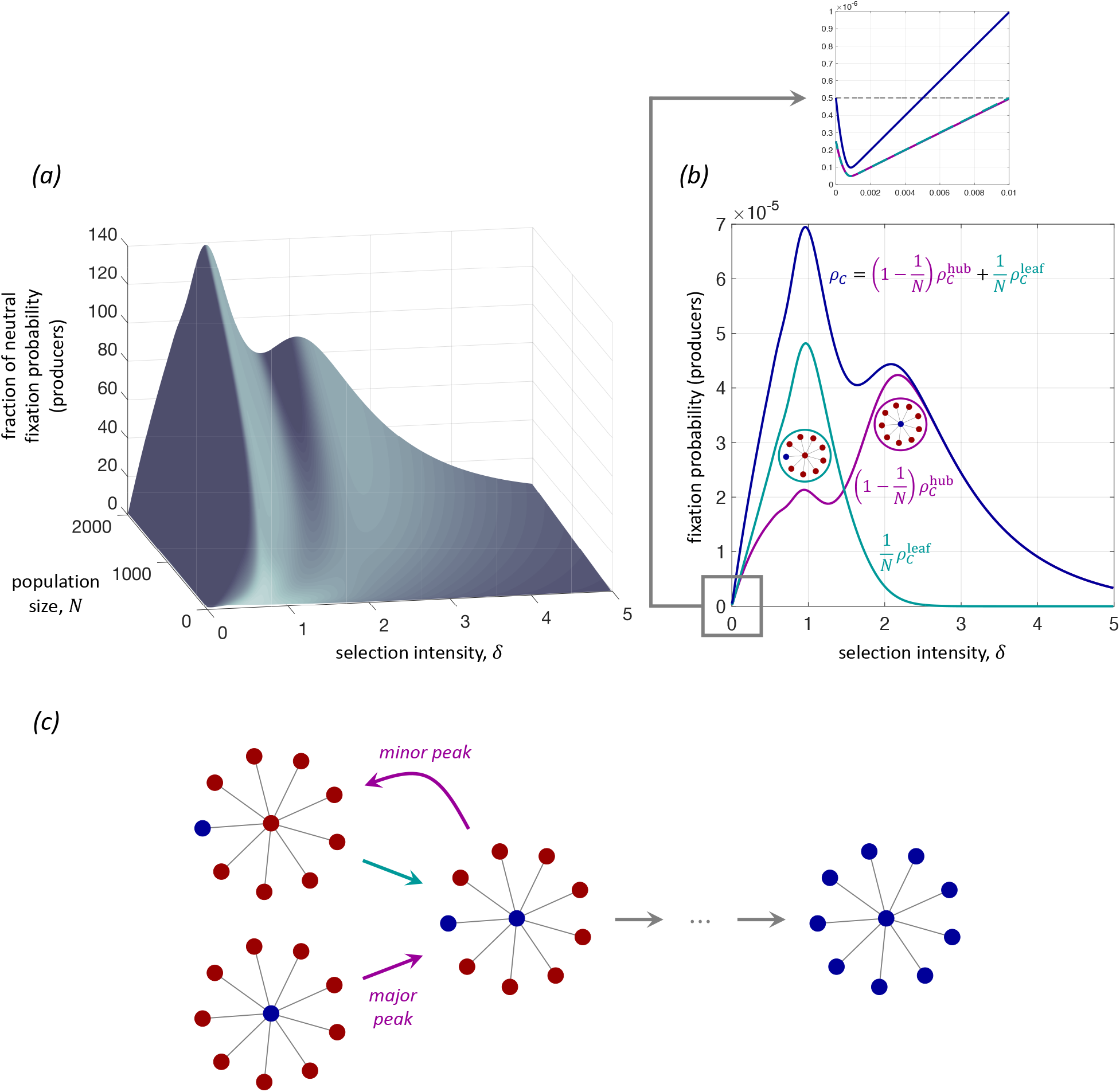
Fixation probability of producers of *ff*-goods on the star, under temperature initialization, as a function of selection intensity, *δ*, and population size, *N* (exact calculations). These curves generalize the blue curve for the fixation probability of producers in **Fig. 4**(*f*). (*a*) Unlike in the case of smaller population sizes, *ρ_C_* can have multiple local maxima when *N* is larger. (*b*) When *N* = 2000, the peak fixation probability of producers is magnified by a factor of nearly 140 over its neutral value. In the inset of (*b*), the black dashed line represents the fixation probability under neutral drift. For all population sizes depicted here, producers are initially disfavored by weak selection (sufficiently small *δ*; see inset). (*c*) For the star, one peak can be attributed to mutant producers arising at the leaf nodes 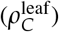, while the other can be attributed to mutant producers arising at the hub 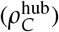. From both configurations, for the mutant type to fix the population must pass through a state with two mutants, one at the hub and one at a leaf node. This state can revert to a configuration with a single mutant at a leaf, even when the initial mutant arises at the hub. In this case, the population is effectively starting from a different configuration, which gives rise to a minor peak corresponding to the value of *δ* that maximizes the probability of fixation from a leaf node. Parameters: *b* = 5 and *c* = 1.

In **Fig. 5**, the peak for the fixation probability of mutants arising at a leaf node comes before that of mutants arising at the hub. For both kinds of initial mutant configurations, the population must transition to a state with a producer at the hub and a single producer at a leaf node on the way to fixation. If the mutant starts at a leaf node, then this producer has payoff – *c* and competes to reproduce with *N* – 2 individuals with payoff 0 and one individual with payoff *b*. If the mutant starts at the hub, then this mutant has payoff –*c* and competes with *N* – 1 individuals with payoff *b* / (*N* – 1). The probability of transitioning to this intermediate two-mutant state is *e*^−*δc*^/ (*e*^−*δc*^ + *e^δb^* + *N* – 2) in the former case and *e*^h*δc*^/ (*e*^−*δc*^ + (*N* – 1) *e*^*δb*/(*N*–1)^ in the latter case. In both cases, this transition is disfavored by selection; but since *e^δb^* – 1 > (*N* – 1) (*e*^*δb*/(*N*–1)^ – 1) for all *N* > 1 and *b*, *δ* > 0 by Bernoulli’s inequality, it is less likely to happen when the mutant arises in a leaf than when it arises in the hub. Therefore, selection must be weaker in the former case than in the latter for the two transitions to have equal probability.

The presence of a “minor” peak in **Fig. 5**(*b*) can be attributed to the fact that, after transitioning from a state with a producer at the hub to one with a producer at the hub and at a leaf, there is a non-zero probability of losing a mutant producer again. After losing a producer, the resulting state has a single producer at a leaf, which represents a different rare-mutant initial configuration (see **Fig. 5**(*c*)). The process then starts in a state for which the optimal fixation probability occurs at a smaller value of *δ*, contributing to a smaller “minor” peak in **Fig. 5**(*b*).

Finally, we turn to the effects of changing the benefit-to-cost ratio of producing a social good. **Fig. 6** shows that increasing the benefit, *b*, while keeping the cost fixed at *c* = 1, tends to move the two peaks at which a producer’s fixation probability is optimized toward smaller values of *δ*. **Fig. 6** also suggests that for fixed selection intensity *δ*, increasing *b* has an effect only until until a certain point. However, we note that although the contour lines on the right-hand side of **Fig. 6** appear approximately vertical, this is merely a consequence of the moderate range of possible values for *b*. The explicit expressions for fixation probabilities given in Methods shows that, for fixed *N* > 1 and *c*, *δ* > 0, the fixation probability of producers approaches zero as *b* → ∞. These expressions elucidate, more generally, the complicated dependence of fixation probabilities on *N*, *b*, *c*, and *δ* when *δ* is neither small nor large, which we have illustrated here numerically.

**Figure 6:**
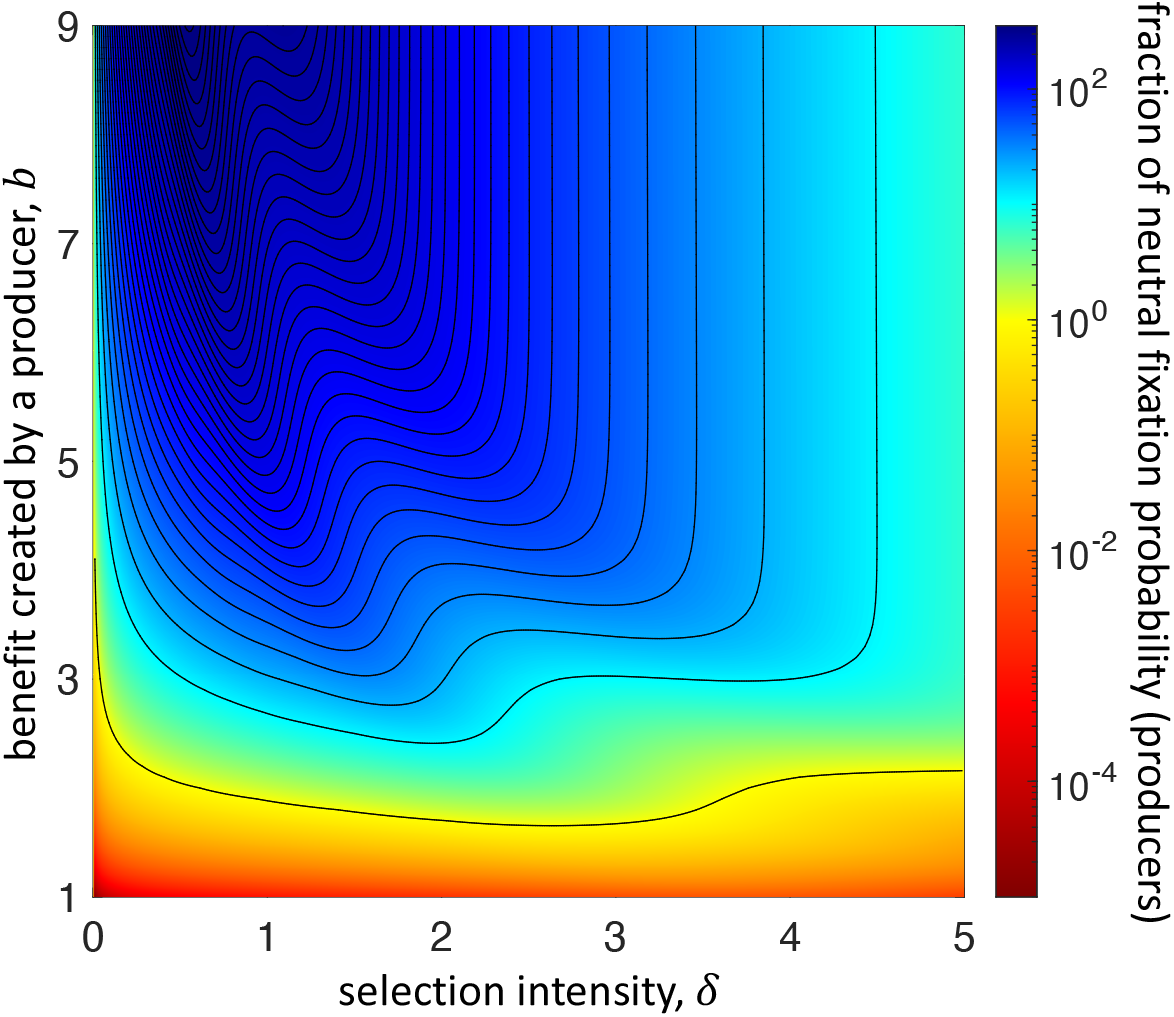
Fixation probability of producers of *ff*-goods on the star, under temperature initialization, as a function of selection intensity, *δ*, and the benefit created by a producer, b (exact calculations). The population size is *N* = 2000. The lowest contour line represents the region for which a producer’s fixation probability is the same as its neutral value. (Note that this neutral value does not depend on *b*.) Neighboring contour lines represent differences of 10 in the ratio of fixation probability to that of neutral drift. As *b* increases, the two peaks highlighted in **Fig. 5** persist, occur at smaller values of *δ*, and increase in magnitude. Notably, for moderately large *δ* beyond these peaks, the contour lines become approximately vertical. For fixed *δ* within this region, increasing *b* initially has little effect on a producer’s fixation probability. However, as *b* grows sufficiently large, this fixation probability does eventually decay to zero.

## 3 Discussion

The four ways in which we have quantified selection favoring type c, namely (*A*^0^), (*A*′), (*R*^0^), and (*R*′), are all long-term measures [28] of evolutionary dynamics. As long as the selection intensity is finite, the population must eventually reach a monomorphic state, so all of the fixation probabilities considered here are well-defined. However, one might argue that the transient dynamics can be more important in models of non-weak selection than in those with weak selection. For example, when there is global competition to reproduce, stronger selection can essentially localize the dynamics for long periods of time. To see why, consider a highly-connected hub within a larger, heterogeneous graph with *Bd* updating. Locally, this hub resembles the star of **Fig. 1**(*b*). If the hub and all of its neighbors are producers, then the hub has an extremely large payoff relative to other individuals in the population. Since competition is global, this hub is involved in all tournaments for reproduction. If at least some other individuals in the population are non-producers, an individual other than the hub must be selected to reproduce for the state to change. But this might happen only rarely, especially when fecundity grows exponentially in payoff and selection intensity. These transient states, which resemble the metastable coexistence states observed in other games [34], are also relevant in addition to the long-term metrics considered here.

The conditions imposed on the payoff-to-fecundity map do not necessarily need to hold in every population. This is especially true of (*v*), which says that shared background payoff/wealth does not affect competition to reproduce. It might be reasonable that fecundity is zero whenever *u* is less than some threshold payoff, in which case competition is not purely relative. Fecundity could also be a concave or linear function of payoff. All of these functional forms depend on the underlying nature of competition for reproduction and/or survival. The assumption that we ignore any background payoff shared by all members of a population is reasonable in light of the focus on just two strategies, *C* and *D*. After all, if *D* is the status quo, which includes all background traits irrelevant to the adoption of a new trait, *C*, and fecundity could be interpreted as being associated to the adoption of *C* only, rather than to the organism as a whole. What is more important in this model is that the exponential payoff-to-fecundity is not arbitrary. It arises as the unique map satisfying a number of simple and intuitive conditions based on the nature of competition. Ideally, other payoff-to-fecundity maps should be derived from similar assumptions and not chosen arbitrarily from some class of maps (e.g. from the class of concave maps).

To our knowledge, our examples provide the first demonstrations that selection can favor the evolution of prosocial traits under Birth-death updating. It is worth mentioning that *Bd* updating and weak selection can support cooperation in weaker social dilemmas. For example, in the snowdrift game, spatial structure can increase the equilibrium frequency of cooperation from that of an unstructured population, and indeed the right choice of payoffs can eliminate defectors altogether [35]. The sharp contrast between *Bd* and *dB* updating seen in the prisoner’s dilemma does not hold for the snowdrift game, which may be attributed (in part) to the fact that defection is not a dominant action in this game [11]. In more stringent conflicts of interest like the production of social goods, where defection is dominant, non-weak selection can draw richer behavior out of social dilemmas, even resulting in the proliferation of altruistic traits when there is global competition to reproduce.

This proliferation of producers, which is a result of a “sweet spot” for the balance of drift and selection, is closely related to how heterogeneous populations generate wealth stratification [36]. While *pp*-goods can result in an unequal distribution of payoffs, this inequality is significantly amplified by *ff*-goods [20]. On the star, for precisely this reason, states that are favorable to the hub for *pp*-goods are even more favorable for *ff*-goods. As a result, the overlap between the selection intensities resulting in the appearance of favorable configurations and the selection intensities driving those configurations toward the all-producer state is larger for *ff*-goods than it is for *pp*-goods. In fact, there is no such overlap at all for *pp*-goods, as shown in **Fig. 4**(*a*,*d*).

Our use of exact calculations in small populations with simple spatial topologies is motivated by two factors. First, there are technical obstacles, computationally, to determining whether selection favors cooperation if the intensity of selection is not sufficiently small [37]. Second, one way of accounting for payoffs here (concentrated benefits) involves stochastic payoffs. These are considerably more difficult to analyze directly, although they are perhaps even more relevant than deterministic payoffs in real populations. And, importantly, they cannot be modeled using expected (mean) payoffs unless one can prove that such a model results in the same dynamics. More than anything, both the update rules and the population structures should be understood as simple examples to illustrate what we believe is important behavior that can arise in the dynamics of models with non-weak selection. We anticipate that teasing out the effects of non-weak selection in other heterogeneous populations will be difficult, but that ultimately these kinds of models will exhibit much more fascinating behavior than those with extremely weak or strong selection. The patterns explored here through the relevant abstractions of the cycle (or regular graphs more generally) and the star provide important reference points for the dynamics in general networks.

## 4 Methods

### 4.1 Payoff-to-fecundity map

Here, we show that under the five assumptions, (*i*)–(*v*), on the payoff-to-fecundity map, it must be true that *F* (*u*) = *e*^*αu*+*β*^ for some *α* ⩾ 0 and 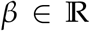. To see why, suppose first that the group in which competition for reproduction takes place is *I* = {1,2}. In this case, condition (*v*) is equivalent to the equation

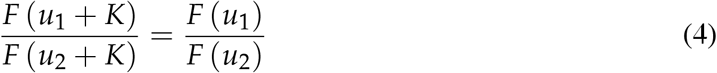

holding for every *u*_1_, *u*_2_, 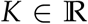. We note that condition (*i*), which says that *F* (*u*) ⩾ 0 for every 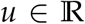, implies that *F* (*u*) > 0 for every 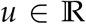 when combined with condition (*v*). Otherwise, competition for reproduction, defined by condition (*v*), would not be well-defined. Letting *u*_2_ = 0, we then have *F* (*u*_1_ + *K*) = *F* (*u*_1_) *F* (*K*) /*F* (0) for every *u*_1_, 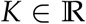. Since *F* is also continuous by (*iii*), it is well-known that this functional equation implies that *F* (*u*) = *e*^*αu*^/*F* (0) for some 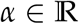 [38]. *F* being a non-decreasing function of *u*, condition (*ii*), is equivalent to *α* ⩾ 0. Letting *β* = − log *F* (0), we then obtain *F* (*u*) = *e*^*αu*+*β*^. This function clearly also satisfies all of the desired properties for larger sets, *I*, which gives the claimed result.

Since selection intensity scales payoffs, (*iv*), the fecundity attributed to payoff *u* at intensity *δ* is *e*^*αδu*+*β*^. But competition is relative, so the probability that *i* reproduces when competing with the individuals in *I* is *F_i_* / ∑_*j*∈*I*_ *F_j_* = *e*^*αδu_i_*^ / ∑_*j*∈*I*_ *e*^*αδu_j_*^ (also called a “softmax function” [39] or a “Gibbs measure” [40]). Thus, without a loss of generality, we may assume that *β* = 0. As a result, up to a rescaling of the selection intensity by *α*, there is a unique payoff-to-fecundity function satisfying the five assumptions imposed in the text.

If selection acts on survival rather than on reproduction, then the assumptions on the payoff-to-fecundity map can be modified in a straightforward manner. Specifically, if *F* quantifies how likely an individual is to be chosen for death, then the corresponding assumptions are

i. *F* (*u*) ⩾ 0 for every 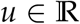;
ii. *F* is non-increasing;
iii. *F* is continuous;
iv. selection intensity scales payoffs, i.e. the survival quantifier associated to payoff *u* at intensity *δ* ⩾ 0 is *F* (*δu*);
v. for every competition within a group *I* ⩾ {1,…, *N*}, the probability that *i* ∈ *I* is chosen for death is invariant under adding a constant, *K*, to the payoffs of all competing individuals in *I*. That is, if *u_i_* and *F_i_* are the payoff and survival quantifier of individual *i*, then

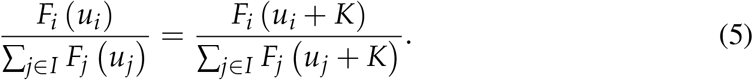

A similar argument then yields *F*_i_/ ∑_*j*∈*I*_ *F_j_* = *e*^−αδu_i_^ / ∑_*j*∈*I*_ *e*^−αδu_j_^, where *α* ⩾ 0.

### 4.2 Calculation of fixation probabilities

Here, we briefly outline the Markov chains defined by the two examples considered in the text, and how they are used to calculate fixation probabilities. Each of these chains is defined on some state space, *S*, and is specified by probabilities *P*_x→y_ for x, y ∈ *S*, representing the probability of transitioning from x to y. Let C and D denote the monomorphic all-producer and all-non-producer states, respectively. The probability of reaching C from state x may be found by numerically solving the recurrence relation *ρ_C_* (x) = ∑_y∈*S*_ *P*_x→y_*ρ_C_* (y), together with the boundary conditions *ρ_C_* (C) = 1 and *ρ_C_* (D) = 0. The probability of reaching the complementary state, D, is then just *ρ_D_* (x) = 1 – *ρ_C_* (x).

#### 4.2.1 Cycle with *dB* updating

Since the cycle is one-dimensional and invading mutants spread in a cluster, the state space for the process may be reduced to *S* = {0,1,…, *N*}, which has size *N* + 1. In particular, a state *n* represents the number of *C* in a cluster, with the remaining *N* – *n* individuals forming a cluster of *D* individuals. Provided *n* is neither 0 nor *N* (the absorbing states), the state can transition from *n* to *n* – 1 or *n* + 1; otherwise, it remains in *n*. Since we allow for stochastic payoffs, we must keep track of the fecundity of each individual. Let *F_i_* (*n*) denote the fecundity of individual *i* in state *n*, which is allowed to be a random variable and correlated with *F_j_* (*n*) for *j* ≠ *i*. To calculate the probability of gaining a mutant in state n under *dB* updating, we first calculate the probability that such a transition happens for a fixed fecundity vector 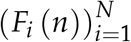. To do so, we only need to examine the boundaries between *C* and *D* on the cycle. Provided 0 < *n* < *N*, there are two such boundaries, although the individuals in these two boundaries overlap when *n* = 1 or *n* = *N* – 1.

As an example, suppose that 1 < *n* < *N* – 1. On one of the two boundaries, suppose that *i* – 1 is a *C* and *i* is a *D*. To gain a *C* along this boundary, increasing the state from *n* to *n* + 1, *i* must first be selected for death, which happens with probability 1/*N*. Then, *i* – 1 (type *C*) must win in a competition for reproduction against *i* + 1 (type *D*). The latter happens with probability *F*_*i*–1_ (*n*) / (*F*_*i*–1_ (*n*) + *F*_*i*+1_ (*n*)). The other boundary, as well as the cases with *n* = 1 or *n* = *N* – 1, are analyzed similarly-except, in the latter case, a single individual is part of both boundaries. Finally, to get the overall transition probabilities, we take expectations over the transition probabilities that were calculated for fixed 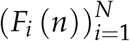, where the source of randomness is the stochasticity in the payoffs that lead to fecundity.

#### 4.2.2 Star with *Bd* updating

Calculations on the star are done similarly, except that in this case the state space is *S* = {0,1} × {0,1,…,*N* – 1}, which has size 2*N*. An element (*m*,*n*) in this space represents whether the individual at the hub is a producer (*m* = 1) or a non-producer (*m* = 0), as well as how many individuals of type *C* are among the leaves, *n*. Under *Bd* updating, the probability that *i* reproduces in state (*m*, *n*) is 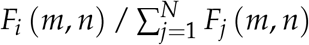, and the offspring replaces a randomly-chosen neighbor. As before, we are interested in the expected value of this probability, taken with respect to the distribution over 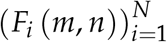 due to payoff stochasticity. Our calculations are based on numerical solutions to the linear systems describing fixation probabilities.

More explicit expressions for fixation probabilities on a star graph have been derived by Hadjichrysathou et al. [41], which reveal a complicated relationship among *N*, *b*, *c*, and *δ*. If *ρ_C_* (*i*, *j*) is the probability of hitting (1, *N* – 1) (the all-producer state) when starting from (*i*, *j*), then for the two kinds of rare-mutant states on the star, Hadjichrysathou et al. [41] show that if 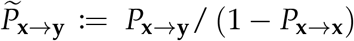 is the probability of transitioning from x to y, conditioned on the system not staying in state x (a non-absorbing state), then

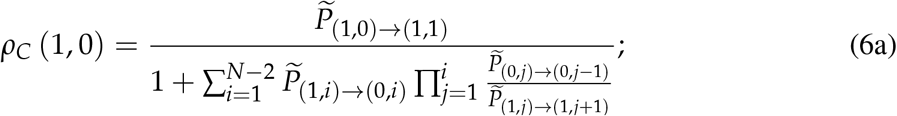

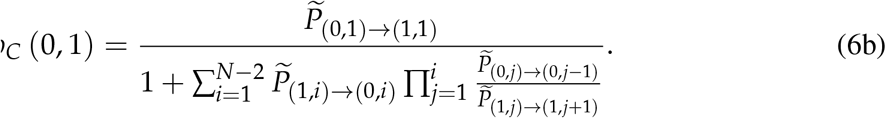

For *ff*-goods, which was not considered by Hadjichrysathou et al. [41], we have

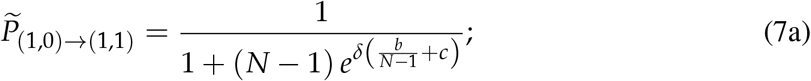

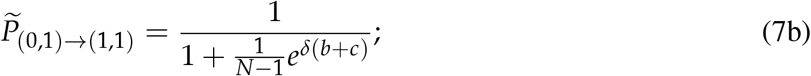

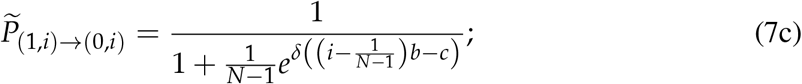

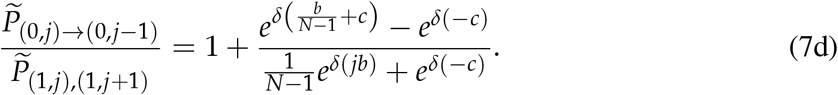

Unlike the case of weak selection, where one can linearize the expressions in **Eq. 6** around *δ* = 0, the balance of drift and selection is much more complicated for larger *δ*. On the star, **Eqs. 6–7** give the qualitative dependence of the long-term success of producers of *ff*-goods as a function of population size, benefit, cost, and selection intensity.

## Acknowledgments

We are grateful to Christian Hilbe, Yoichiro Mori, Joshua Plotkin, and Qi Su for helpful conversations. A.M. was supported by the Simons Foundation (Math+X Grant to the University of Pennsylvania). C.H. acknowledges funding by the Natural Sciences and Engineering Research Council of Canada (NSERC), grant RGPIN-2015-05795. The funders had no role in study design, data collection and analysis, decision to publish, or preparation of the manuscript.

## Data availability statement

Supporting code is available at https://github.com/alexmcavoy/intermediate-selection.

